# Harnessing Colon Chip technology to identify commensal bacteria that promote host tolerance to infection

**DOI:** 10.1101/2020.12.05.412353

**Authors:** Francesca S. Gazzaniga, Diogo M. Camacho, Meng Wu, Matheus Palazzo, Alexandre Dinis, Francis N. Grafton, Mark J. Cartwright, Michael Super, Dennis L. Kasper, Donald E. Ingber

## Abstract

Commensal bacteria within the gut microbiome contribute to development of host tolerance to infection, however, identifying specific microbes responsible for this response is difficult. Here we describe methods for developing microfluidic organ-on-a-chip models of small and large intestine lined with epithelial cells isolated from duodenal, jejunal, ileal, or colon organoids derived from wild type or transgenic mice. To focus on host-microbiome interactions, we carried out studies with the mouse Colon Chip and demonstrated that it can support co-culture with living gut microbiome and enable assessment of effects on epithelial adhesion, tight junctions, barrier function, mucus production, and cytokine release. Moreover, infection of the Colon Chips with the pathogenic bacterium, *Salmonella typhimurium*, resulted in epithelial detachment, decreased tight junction staining, and increased release of cytokines (CXCL1, CXCL2, and CCL20) that closely mimicked changes previously seen in mice. Symbiosis between microbiome bacteria and the intestinal epithelium was also recapitulated by populating Colon Chips with complex living mouse or human microbiome. By taking advantage of differences in the composition between complex microbiome samples cultured on each chip using 16s sequencing, we were able to identify *Enterococcus faecium* as a positive contributor to host tolerance, confirming past findings obtained in mouse experiments. Thus, mouse Intestine Chips may represent new experimental *in vitro* platforms for identifying particular bacterial strains that modulate host response to pathogens, as well as for investigating the cellular and molecular basis of host-microbe interactions.

## INTRODUCTION

The epithelium lining the intestine is an essential barrier that protects our bodies from the trillions of microbes within the microbiome that resides within its lumen. The intestinal epithelium regulates this critical barrier by integrating signals from commensal bacteria within the microbiome with cues from immune cells in the lamina propria, such that a state of host tolerance develops which enables host-microbiome symbiosis. Maintaining this state of tolerance in which microbes and the host co-exist without causing damage to each other is essential to protect against a wide variety of diseases. Great advances in our understanding of host-microbiome relations has been made through use of studies in mice where both the composition of the microbiota and host immune cell regulatory responses have been well characterized(1–3). However, it is extremely difficult to identify specific cellular and molecular contributors to host tolerance using *in vivo* experiments alone. Recently developed organ-on-a-chip (Organ Chip) microfluidic culture technology offers a way to recapitulate organ-level structures and functions *in vitro*. Thus, in the present study, we set out to explore if Organ Chip technology could be leveraged to develop an *in vitro* model of the mouse intestinal barrier in the colon that can be used to study host-microbiome interactions that contribute to tolerance development.

Studies in mice suggest that gut bacteria can protect from infection by several mechanisms. For example, differences in gut microbiota contribute to species-specific differences in resistance to *Salmonella typhimurium* infection where the half maximal infectious dose (ID_50_) is more than three logs higher in mice than in humans, unless the mice are pretreated orally with streptomycin to alter the microbiome (4). Furthermore, when gnotobiotic mice that lack a microbiome are recolonized with healthy mouse microbiota, they become more resistant to *S. typhimurium* infection than when colonized with healthy human microbiota (2).

In addition to colonization resistance, the microbiome can also protect from infection by inducing host tolerance. *Enterococcus faecium*, a bacterium found in healthy human stool, promotes tolerance to *S. typhimurium* in mice as well as in *C. elegans* (5,6). *S. typhimurium* colonizes the intestine of mice containing *E. faecium*, but disease severity is limited due to thickening of the mucus layer, which prevents this potential pathogen from accessing the gut epithelium. In other cases, gut bacteria prevent infection by impacting the host immune system. For instance, Type 3 Innate Lymphoid Cells (ILC3s) located in the intestinal lamina propria protect against infection by responding to microbial signals and releasing the cytokine interleukin-22 (IL22), which promotes production of antimicrobial peptides (7). With the rise of antibiotic resistance, understanding how microbes impact the gut epithelium to maintain a tolerance state is critical for developing new and more effective anti-infective therapeutics.

Microfluidic Organ Chips lined by living cells isolated from host tissues can be used to recapitulate organ level behaviors with high fidelity *in vitro*, and to interrogate cellular interactions with great precision in an organ-level context (8–11). Human Intestine Chips have been created that are lined with primary epithelial cells isolated from patient-derived organoids(9,11,12), which provide multiple advantages over organoid culture alone. In contrast to organoids, Organ Chips enable culture of complex living microbiome in direct contact a differentiated intestinal epithelium and its overlying mucus layer while experiencing dynamic flow and peristalsis-like motions, in addition to providing direct access to the apical and basal compartments so that barrier, transport, and absorptive functions can be measured (8,9,13). Individual cell types also can be added or removed in different combinations to Organ Chips to pinpoint the directionality of cell-cell and tissue-tissue interactions. For example, human Colon Chips were recently used to investigate how microbiota metabolites from mouse versus human feces impact susceptibility to enterohemorrhagic *E. coli* infection (13). This system replicated species-specific findings that mouse microbial metabolites provided better protection from infection than those from human microbiome, and led to the identification of human microbiome metabolites that increase susceptibility to infection (13).

While most Organ Chips have used human cells to mimic human physiology (10,14), chips created with mouse intestinal epithelial cells would provide the benefit of being able to directly compare chip responses to results of prior animal studies, while also providing the ability to isolate and add physiologically relevant cells that are often difficult to obtain from human biopsies. Here we describe methods for developing mouse Intestine Chips lined by epithelial cells isolated from duodenum, ileum, jejunum or colon. We also show that the Colon Chip supports co-culture of complex gut microbiome with differentiated intestinal epithelium and show that this experimental model can be used to investigate specific host-microbe interactions under controlled conditions *in vitro*.

## Materials and Methods

### Isolation of mouse intestinal crypts

C57/Bl6 WT or Kaede mice were sacrificed by CO_2_ asphyxiation followed by cervical dislocation and the large and small intestine was removed. The small intestine was separated into three segments: duodenum, jejunum, ileum and colon. All intestinal crypts were isolated based on the Stem Cell Technologies protocol. Intestinal segments were flushed with cold PBS and cut lengthwise and then cut into 2 mm pieces and placed into a 50 ml conical tube with 10 ml cold PBS. Using a pre-wetted 10 ml pipette, intestine pieces were pipetted vigorously up and down three time. After intestinal pieces settled to the bottom of the tube, the supernatant was removed. This procedure was repeated until the supernatant was clear (10-20 times). Clear supernatant was removed and intestinal pieces were resuspended in 25 ml of Gentle Cell Dissociation Reagent (Stem Cell Technologies, 07174). Tubes were incubated at room temperature on a rocking platform for 20 minutes. Tubes were removed from the rocking platform and intestinal pieces were left to settle to the bottom of the tube. Once the pieces settled, the supernatant was removed. Intestinal pieces were vigorously pipetted up and down three times in 10 ml of cold PBS + 0.1% BSA and left to settle on the bottom of the tube. Supernatant was collected, passed through a 70 micron filter, and labeled as fraction 1. This step was repeated for a total of 4 fractions. Fractions were spun down at 290 g for 5 minutes at 4°C. Supernatant was removed and fractions were resuspended in 10 ml cold PBS + 0.1% BSA. 1 ml of each fraction was placed in a 24 well dish and the quality of the crypts were assessed by an inverted microscope. The fraction with the most crypts and fewest single cells was chosen. Crypts were counted and spun down 290 g for 5 minutes at 4°C. Crypts were resuspended in growth factor reduced Matrigel (Corning, 356231) at 30 crypts per microliter. 50 microliters were plated onto 24 well non-tissue culture treated plates (Costar 3738). Plates were placed in 37 °C incubator for 15 minutes to allow for Matrigel to solidify. 500 microliters of mouse organoid culture media (Advanced DMEM/F12 (12634028 Thermo Fisher Scientific), 50% L-WRN (Wnt3a, R-spondin, Noggin) conditioned media(16), 2 mM glutamax (35050061 Thermo Fisher Scientific), 10 mM HEPES (15630080 Thermo Fisher Scientific), 50 ng/ml murine epidermal growth factor (Invitrogen PMG8043), N2 supplement (Invitrogen, 17502-048) B27 supplement (Invitrogen 17504-044), 1 mM N-Acetylcystine (Sigma Aldrich A9165-5G), 100 μg/ml Primocin (Invivogen ant-pm-1)) plus 10 uM Y-27632 (Sigma Aldrich Y0503) was placed on top of the Matrigel domes. Media was changed three times a week and Y27632 was only added the day of splitting.

### Mouse intestine chip cultures

The mouse colon chips use the same chip design, membrane activation, membrane coating, organoid harvesting and seeding as previously described (9,12,14,27). Briefly, microfluidic chips composed of PDMS and two parallel microchannels separated by a porous membrane were purchased from Emulate, Inc (CHIP-S1 Stretchable Chip, RE00001024 Basic Research Kit; Emulate, Inc). The inner surfaces of both channels were activated with 0.5 mg/ml sulfo-SANPAH solution (A35395, Thermo Fisher Scientific) as in (27). The inner surfaces of both channels and membrane were coated with 200 ug/ml rat tail collagen type I (354236, Corning) and 4% Matrigel (Corning, 356231) as previously described (9). Mouse intestine organoids were maintained in 50 μl of Matrigel in a 24 well dish with 500 ul mouse organoid media and split at 1:4 ratio once per week. Two days before chip seeding, organoids were split as previously described (12). Organoids were isolated from Matrigel and fragmented as previously described (12). Organoids were resuspended in mouse organoid medium supplemented with 10 μM/L Y-27632 at 6 x 10^6^ cells/ml. The basal channel of extracellular matrix coated chips was filled with 50 μl of mouse organoid media supplemented with 10 μM/L Y-27632 and plugged with 200 ul filter tips at the outlet and inlet. 35 μl of disrupted organoids were seeded onto the top channel and the outlet and inlets were plugged with 200 ul filter tips. Chips were incubated overnight at 37C in 5% CO_2_ to promote cell adhesion. The following day, the apical channel was washed with 100 ul of mouse organoid media at a time until all unattached cells were removed. The chips were placed in the anaerobic farms and connected to peristaltic pumps as previously described (14). Chips were perfused with mouse organoid media on the apical and basal channels at 1 μl/ minute. Media was added to reservoirs every other day. Mouse intestine chips formed confluent monolayers one week after culture. After monolayer formation, anaerobic gas (5% CO_2_ 95 % N_2_) was perfused through the apical channel of the anaerobic farm as previously described (14).

### Immunofluorescent Microscopy

Both channels of mouse colon chips were washed with 100 μl of PBS and then fixed with 50 μl of 4% PFA for 15 minutes at room temperature. Chips were washed with 100 μl of PBS and blocked and permeabilized with 100 μl of 0.1% Triton X-100 and 5% BSA in PBS for 1 hour at room temperature. Samples were stained overnight with the following primary antibodies diluted 1:100 in 5% BSA in PBS at 4C: anti-MUC2 (Santa Cruz Biotechnology sc-15334), anti-ZO-1 (Santa Cruz Biotechnology sc-33725), anti-ChrA (Santa Cruz Biotechnology sc-1488), anti-phalloidin 647 (Thermo-Fisher A22287). The following day, chips were washed with 100 ul PBS per channel and stained with the following secondary antibodies diluted 1: 100 in 5% BSA in PBS for 2 hours at room temperature: goat anti-rat Alexa Fluor 488 (Thermo Fisher Scientific A11006), donkey anti-rabbit Alexa Fluor 647 (Thermo Fisher Scientific A31573), donkey anti-goat Alexa Fluor 555 (Thermo Fisher Scientific A21432). Chips were washed with 100 ul of PBS three times. During the second wash, chips were incubated with 5 μg/ml DAPI (Thermo Fishcer Scientific D1306) for 5 minutes at room temperature. Chips were imaged with a laser scanning confocal microscope (Leica SP5 X MP DMI-6000). Images were processed in FIJI2 and nuclei coverage and ZO-1 intensity was analyzed using FIJI2. For imaging Kaede chips, Kaede chips were photoconverted by shining a 405 nm laser for 30 seconds.

### Cytokines/chemokines analysis

Cytokine and chemokine levels were measured in the basal outflow of mouse intestine chips using a custom MSD U-plex Assay (Meso Scale Diognostics) and the mouse CXCL1/KC DuoSet ELISA (R&D Systems DY453) according to the manufacturers’ instructions.

### Generation and culture of Salmonella typhimurium mCherry

To generate *S. typhimurium-mCherry*, the plasmid pAW83-mCherry was isolated with a miniprep kit (Qiagen) and then transfected into electrocompetent *Salmonella enterica serovar typhimurium* strain SL1344 through the use of electroporation. Transfected cells were grown on selective LB agar with 100 μg/mL carbenicillin, and a single colony was selected to produce 15% glycerol stocks for −80°C storage as well as confirmation of mCherry expression.

To establish long-lived quantified stocks for infection, the *S. typhimurium-mCherry* was grown to exponential phase in a large volume of LB broth with 100 μg/mL carbenicillin. The exponential growth was then centrifuged, the pellet was washed once, and then reconstituted in ice cold sterile saline/dextrose (5% dextrose and 0.45% sodium chloride) with 15% glycerol to generate an optical density (OD) value equivalent to 1e9 CFU/mL. It was then split across several small aliquots stored at −80°C, and an aliquot was plated after freeze-thaw to confirm CFU accuracy.

### *E. faecium* culture

*E. faecium* isolated from Hmb stock was grown overnight at 37C in aerobic conditions in brain heart infusion media (B11059 BD Biosciences). *E. faecium* was pelleted at 5000g, washed once in DMEM and resuspended in antibiotic free bacterial mouse organoid media (mouse organoid media + 1 mg/ml pectin, 1 mg/ml mucin, 5 μg/ml Hemin, 0.5 μg/ml Vitamin K1) at 1×10^6^ CFU/ml.

### Bacterial colonization of chips

Twenty-four hours before bacterial colonization, reservoirs were washed with PBS and antibiotic free mouse organoid media and antibiotic free bacterial mouse organoid media was added to the basal and apical channels, respectively. On the day of colonization, anaerobic farms were moved to an anaerobic chamber. Basal channels were plugged with 200 ul filter tips. 50 μl of 1×10^6^ *S. typhimurium* or *E. faecium*, or 50 μl of 1:100 Hmb or Mmb stock (generated as in (14)) was seeded into the apical channel. Anaerobic farms were reconnected to pumps and left static for 30 minutes before media was perfused at 1 μl/minute. In some experiments, *E. faecium*, Hmb, or Mmb, was perfused in colon chips for 16 hours before infection with *S. typhimurium*. Chips were flushed at 50 μl/min for two minutes and outflow was plated in eight 10-fold serial dilutions onto Bile Esculin Agar and Xylose Lysine Deoxycholate plates for *E. faecium* and *S. typhimurium* samples or frozen at −80C for 16S sequencing.

### 16S rRNA profiling of the microbiota

Frozen chip outflow was either sent to Diversigen for 16S sequencing (Figure 3A and Figure S6) and analysis (Figure 3A) or processed and analyzed in house using the same primers as Diversigen (Figures 4A,B). When processed in house, bacterial DNA was extracted using QIAamp Fast DNA Stool Mini Kit (Qiagen 51604). Purified DNA was quantified by Qubit dsDNA HS Assay (Invitrogen) and normalized. The V4 region of 16S rRNA gene was amplified with primers 515F and 806R under the previously described conditions (28), and ∼390-bp amplicons were purified and quantified by Qubit dsDNA HS Assay and combined with equal mass to make a pooled library. The pooled library was then subjected to multiplex sequencing (Illumina MiSeq, 251 nt x 2 pair-end reads with 12 nt index reads). Raw sequencing data were processed with QIIME2 pipelines (28). In brief, raw sequencing data were imported to QIIME2 and demultiplexed, then DADA2 were used for sequence quality control and feature table construction. The feature table were further used for alpha and beta diversity analysis, as well as taxonomic analysis.

**Figure 1.**
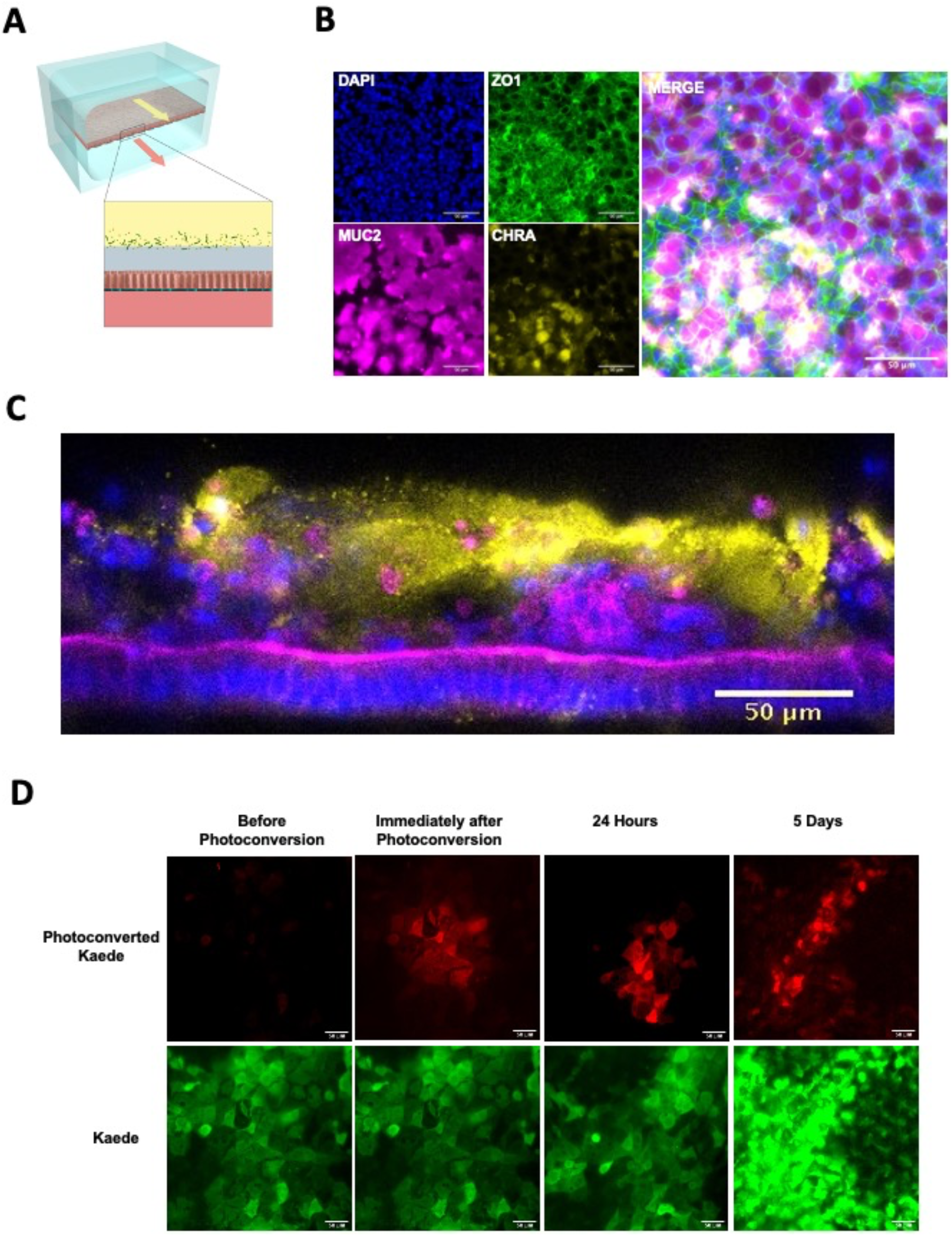
Microfluidic mouse Colon Chips form tight junctions, multiple cell types, and a mucus layer. (**A**) Diagram of a two-channel microfluidic Organ Chip with epithelial cells on top of a porous membrane separating the channels, and bacteria are cultured above the epithelium under a hypoxia gradient that is created by flowing oxygenated medium through the basal channel while maintaining the entire chip in an anaerobic chamber filled with carbon dioxide and nitrogen gas. (**B**) Immunofluorescence microscopic images showing mouse Colon Chip lined by an intestinal epithelium containing tight junctions (ZO-1, green) and goblet cells (MUC2, magenta), and enteroendocrine cells (CHR A, yellow); nuclei are stained with DAPI (blue). Images taken with 25x objective. (**C**) Cross section of mouse Colon Chip showing the polarized epithelium stained for F-actin (magenta) that concentrates at the apical brush border and for MUC2 (yellow), which appears in the overlying mucus layer; nuclei are stained with DAPI (blue). (**D**) Immunofluorescence top-down view showing that mouse Colon Chips form monolayers with cells isolated from organoids derived from Kaede mouse (green), which can be used to track cell divisions. A section of the chip was photoconverted with 405 nm light (red) and imaged daily. Dilution of red signal to green indicates cell turnover over time. Images taken with 25x objective.

**Figure 2.**
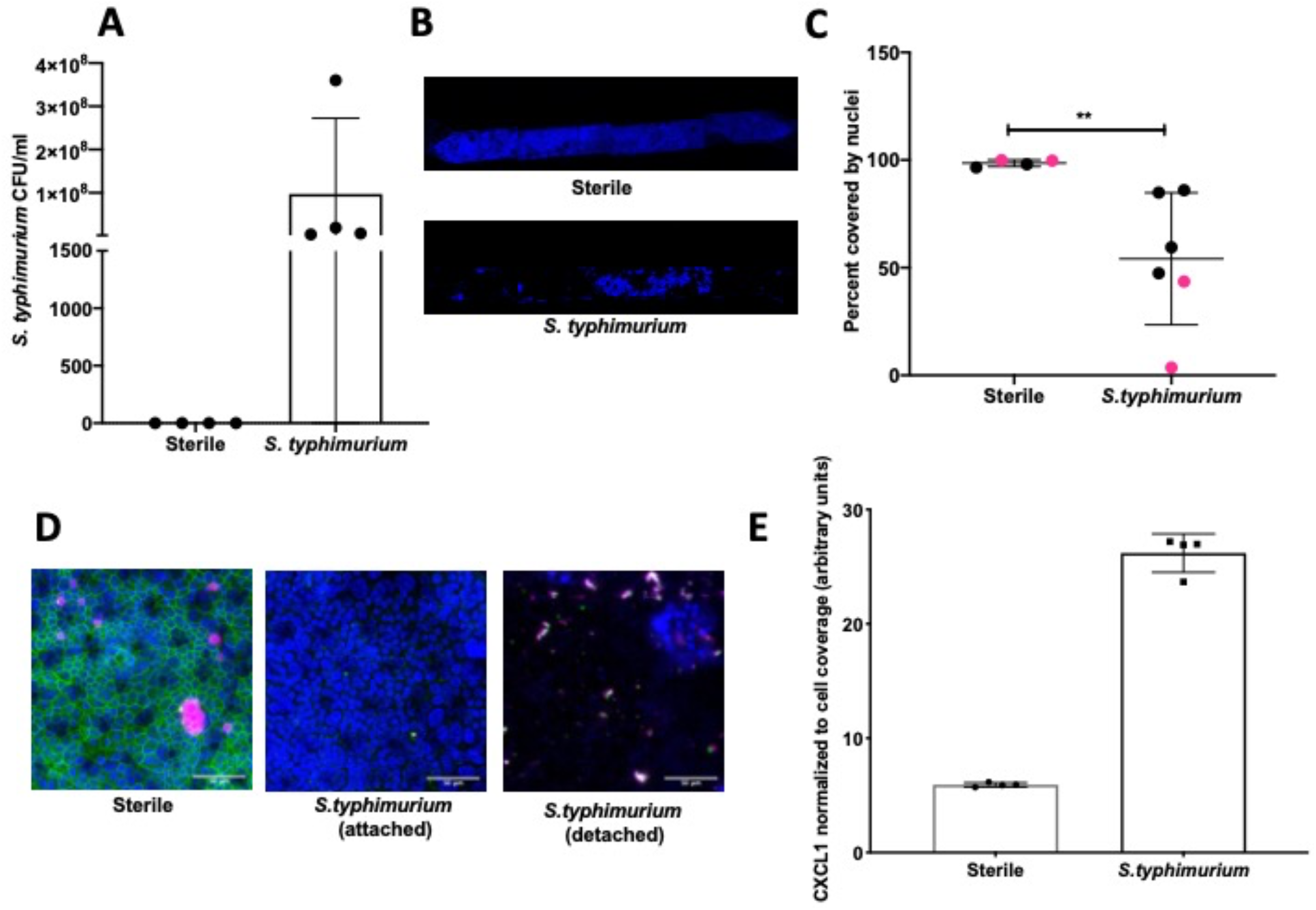
*S. typhimurium* induces epithelial damage in mouse Colon Chips. Chips were inoculated with 5 x 10^5^ CFU/ml and perfused for 24 hours. (**A**) *S. typhimurium* CFU/ml was quantified by plating a fast flush (60 microliters/min) from chips 24 hours after infection onto XLD plates and only detected in chips that were inoculated with *S. typhimurium*. Each dot represents a different chip; similar results were obtained in four different experiments. (**B**) Epithelial lesions induced by *S. typhimurium* infection were visualized by imaging chips in the DAPI channel at 5x, representative examples for Sterile and *S. typhimurium* chips are shown. (**C**) Cell coverage was quantified by processing images using FIJI and calculating the area of the membrane surface covered by cells (shown as percent covered by nuclei). *S. typhimurium* induced significant epithelial cell detachment (Mann Whitney T-test P= 0.0095); data shown are from two representative experiments. (**D**) Sterile and *S. typhimurium*-infected chips stained for DAPI (blue), ZO-1 (green), and MUC2 (magenta). Merged images indicate that *S. typhimurium* disrupts ZO-1 tight junction and reduces MUC2 expression before cell detachment occurs. (**E**) Graph showing CXCL1 levels in outflows of sterile chips versus *S. typhimurium* chips at 24 hours after infection. CXCL1 values were normalized to the membrane surface area covered by cells; similar results were obtained in four different experiments.

**Figure 3.**
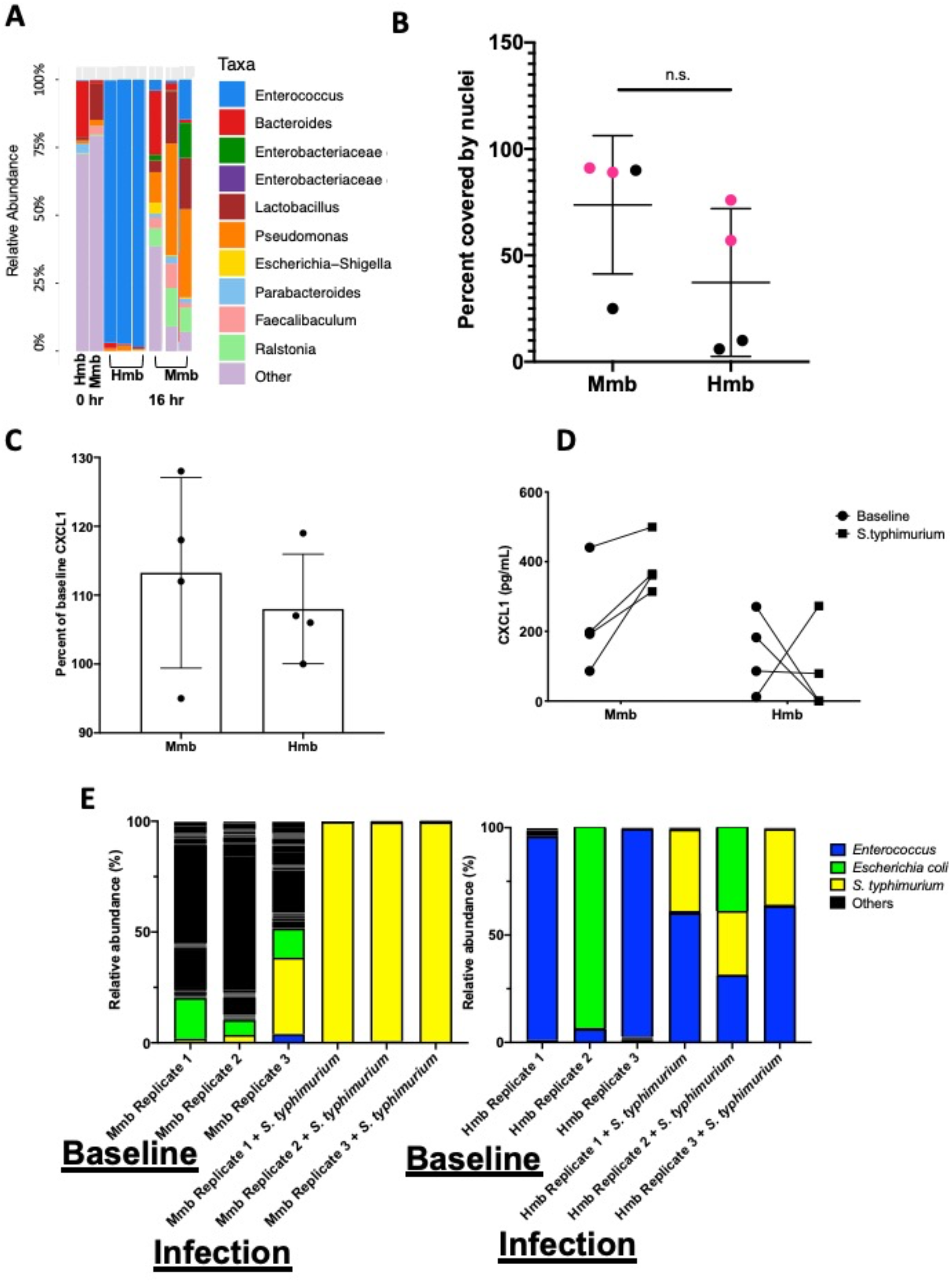
Colon Chips with microbiota can be used to select strains that protect against pathogen. (**A**) Microbiota derived from Hmb or Mmb mice was seeded onto mouse Colon Chips and 16S analysis was performed on seeding stocks and flushes after 16 hours on chip. Relative abundance of the indicated genera reveals *Enterococcus* dominated the Hmb chips. (**B**) Culture surface areas covered by adherent nucleated cells varied between experiments and were not significantly different between Hmb or Mmb colonized chips. Data shown are from 2 different experiments (magenta versus black). (**C**) CXCL1 detected in basal outflows immediately before (baseline) or 16 hours after colonization with Mmb or Hmb. Percent of baseline CXCL1 release shows no significant difference between colonization with Mmb versus Hmb; similar results were observed in four different experiments. **(D)** CXCL1 detected in basal outflows 16 hours after colonization with Hmb or Mmb (baseline; solid circle), and 24 hours after infection with *S*.*typhimurium* (*S*.*typhimurium;* solid square). Hmb colonization, but not Mmb colonization inhibited *S. typhimurium*-induced CXCL1 release. (**E)** 16S sequencing of bacteria in apical outflows collected 16 hours after colonization with Mmb or Hmb (baseline) and 24 hours after infection with *S. typhimurium* (infection) from three replicate chips. Analysis of the relative abundance of genera on chips infected with *S. typhimurium* for 24 hours revealed *Enterococcus* from Hmb correlates with less *S. typhimurium* overgrowth.

**Figure 4.**
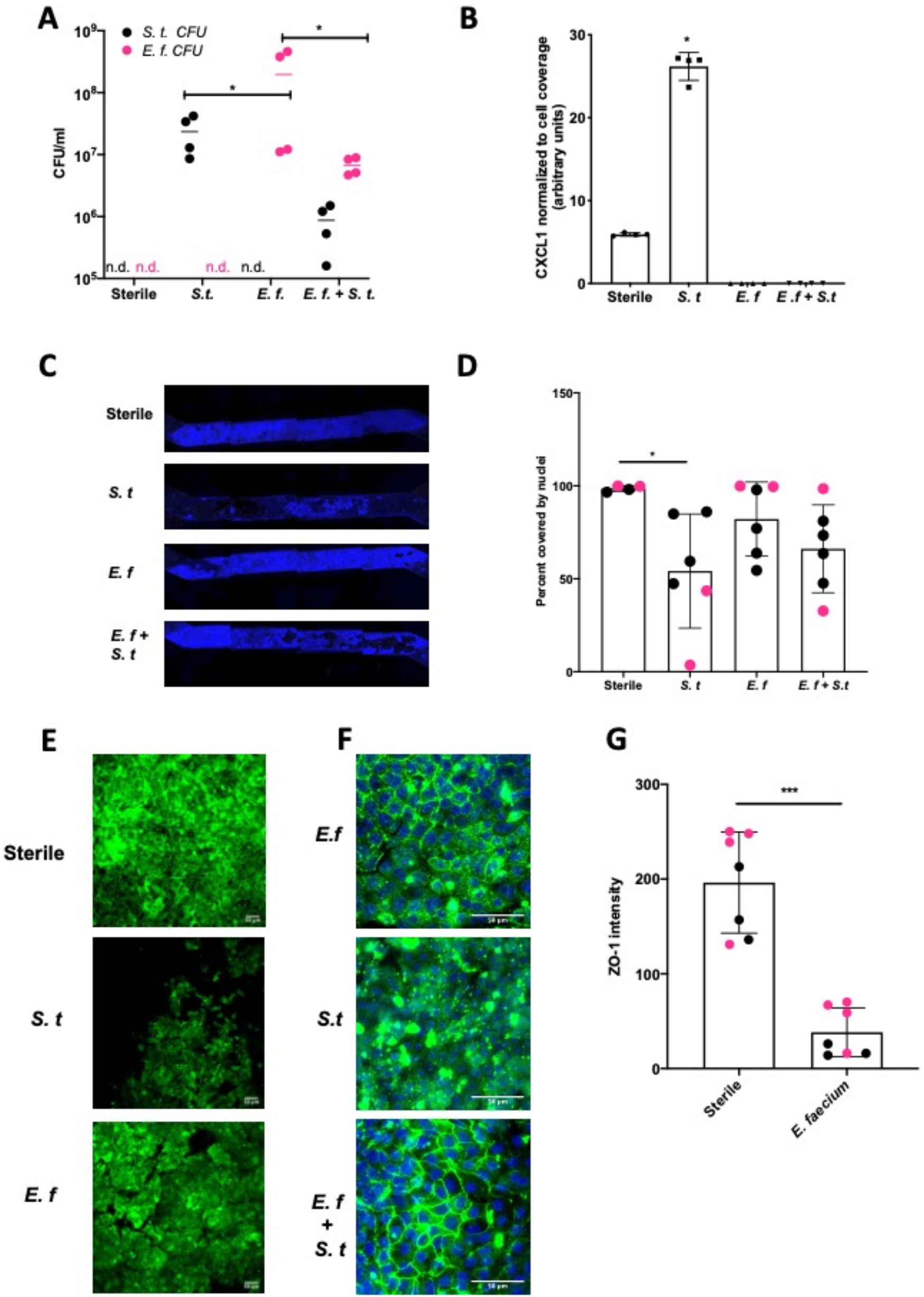
*Enterococcus faecium* protects colon epithelium from *S. typhimurium*-induced damage. Mouse Colon Chips were colonized with *E. faecium* for 16 hours and infected with *S. typhimurium* 24 hours. (**A**). Chips were flushed for 2 minutes and flush was plated in serial dilutions on bile esculin agar (BEA) plates and xylose lysine deoxycholate (XLD) plates to selectively measure *E. faecium* (BEA, magenta) and *S. typhimurium* (XLD, black) growth. Both *E. faecium* and *S. typhimurium* grew when cultured alone in the Colon Chips, but their numbers were reduced by 1 log when both bacteria were cultured simultaneously (similar results were obtained in 4 different experiments). (**B**) Basal chip outflow was collected 24 hours after infection with *S. typhimurium* and CXCL1 measured by ELISA and normalized to cell coverage. Representative example from 4 different experiments. (**C**) Epithelial layer integrity was visualized by imaging DAPI-stained chips at low magnification (5x); representative examples for Sterile, *S. typhimurium, E. faecium, and E. faecium + S*.*typhimurium* chips are shown. **(D)** Cell coverage of the culture surface was quantified by processing images using FIJI and calculated as percent surface area covered by nuclei. *S. typhimurium*, but not *E. faecium* induced significant epithelial cell detachment (adjusted p= 0.0365 by Kruskal-Wallis and Dunn’s multiple comparisons test). Magenta and black dots indicate chips from two different experiments. **(E)** Immunofluorescence images show significant epithelial lesions in *S. typhimurium* infected chips and mild, but not significant epithelial lesions in *E. faecium* chips when compared to sterile chips with completely intact ZO-1 tight junctions (green). Sterile image processed at different exposure from colonized chips (**E**) Representative immunofluorescent images of *E. faecium* monocolonized, *S. typhimurium* monocolonized, and *E. faecium + S. typhimurium* chips show *E. faecium* colonized chips maintain ZO-1 tight junction staining and protects from *S. typhimurium*-reduction of ZO-1 tight junction staining ZO-1 (green), DAPI (blue). (**G**) Quantification of ZO-1 intensity in immunofluorescent images from sterile versus *E. faecium* colonized chips. Mean ZO-1 intensity was measured using FIJI. Combination of two experiments (magenta versus black). Significance determined by Mann-Whitney test. P=0.0006.

## RESULTS

### Mouse Intestine Chips can be generated from wild type or transgenic mice

To develop mouse Intestine Chips, we generated organoids from the colon of wild type (WT) C57/Bl6 mice or transgenic Kaede mice that express a photoconvertible GFP driven by the actin promoter that can be used to track cell divisions and migration when activated with 405 nm light (15). Organoids that were prepared as previously described (16–18) were enzymatically disrupted and physically dissociated before being seeded on the upper surface of the extracellular matrix (ECM)-coated porous membrane that separates the two parallel microfluidic channels of the Organ Chip (**Fig. 1A**). Organoid culture medium was perfused through both channels and a confluent monolayer was observed covering the porous membrane by one week of culture in both WT and Kaede chips, and similar results were obtained when we used intestinal epithelial cells isolated from organoids derived from mouse duodenum, jejunum, and ileum (**Supplementary Fig. S1A-D**).

To study host-microbiome interactions, we focused on Colon Chips. Immunofluorescence microscopic analysis confirmed that the epithelial monolayer forms a continuous network of ZO-1-containing tight junctions and contains MUC2-positive goblet cells as well as enteroendocrine cells that stained positively for chromogranin A staining when viewed from above (**Fig. 1B**). Z stack visualization (**Supplementary Movie S2**) and immunostaining of vertical cross sections through the chips (**Fig. 1C**) revealed a polarized epithelium with basal nuclei, apical junctions enriched for F-actin, and MUC2-staining droplets in the apical regions of the cells as well as a MUC2 rich mucus layer overlying the apical surfaces of the cells. Shed epithelial cells also could be detected in the apical mucus layer suggesting that mouse Colon Chips are able to maintain and replenish the epithelial monolayer as observed in mouse colon in vivo.

To investigate if the intestinal epithelial cells are undergoing active turnover on-chip, sections of chips lined with the photosensitive GFP-expressing Kaede mouse cells were photoconverted and imaged daily. The dilution in the red signal over time within an otherwise continuous epithelium indicated the presence of constant epithelial turnover (**Fig. 1D**). Additionally, qPCR of Colon Chips and organoids after 8 days of culture confirmed the expression of genes encoding the stem cell marker *Lgr5*, and the differentiation markers, *Muc2* and *ChrA* in all tissues (**Supplementary Fig. S3**); however, there was significantly higher levels of *Muc2* and *ChrA* expression in the Colon Chips compared to the organoids. Although *Lgr5* was detectable in both chips and organoids, it was higher in organoids confirming previous reports that the microfluidic chips that experience fluid flow more efficiently promote intestinal differentiation (9,12).

### *S. typhimurium* induces epithelial injury in Mouse Colon Chips

We then assessed if the mouse Colon Chips can be used to interrogate response to pathogens by comparing the responses of the epithelium to co-culture with a known mouse intestinal pathogen, *S. typhimurium* (13-15), versus commensal bacteria from healthy mice. After being seeded in the apical channel of Colon Chips lined by confluent epithelium, mCherry-labeled *S. typhimurium* bacteria grew rapidly as visualized in fixed chips and by live cell imaging (**Supplementary Fig. S4A,B**). Quantification of the bacteria revealed that they increased in number by three logs within 24 hours on-chip (**Fig. 2A**), which is the same rate at which these bacteria grown when administered to germ free mice (2). Within 24 hours of *S. typhimurium* infection, epithelial cell detachment could be detected by a decrease in the chip membrane surface area covered by nucleated cells (**Fig. 2B,C**). When chips were harvested prior to epithelial detachment, ZO-1 and MUC2 staining were reduced (**Fig. 2D**) indicating that disruption of tight junctions and decreased mucus accumulation precede epithelial lesion formation.

To determine how cytokine production in colon chips is impacted by *S. typhimurium* infection, the effluent of the basal channel of the chip was collected at 1.5, 3, 6, and 24 hours after infection and assayed with a multiplex ELISA. We found that CXCL1 and CXCL2 (mouse homologs to IL8) increased at 24 hours following *S. typhimurium* infection, whereas other cytokines were detected at much lower levels (**Supplementary Fig. S5A-C**). Because secretion of CXCL1 & 2 by the epithelium serves to recruit immune cells to the site of infection, we measured CXCL1 in future studies to assess whether the bacteria present within the engineered intestinal lumen would be sensed as an infection requiring induction of an immune response. These subsequent studies confirmed that *S. typhimurium* infection induced more than a 5-fold increase in CXCL1 production in the Colon Chip (**Fig. 2E**). RNA Seq analysis also confirmed that 24 genes were differentially expressed within 6 hours after infection of the Colon Chips with *S. typhimurium*, with almost all (23) genes showing an increase in expression (**Supplementary Fig. S6**). Importantly, all of the genes that were over expressed by the intestinal epithelium in response to *S. typhimurium* infection on-chip (CXCL1, CXCL2, CCL20, and AW112010) behave similarly when mice are exposed to *S. typhimurium* in vivo (19–21). These results indicate that mouse Colon Chips provide a physiologically relevant system to investigate *S. typhimurium* infection in vitro, and that epithelial detachment, decreased tight junction staining, and increased CXCL1 release may be used as metrics to assess bacterial pathogenicity on-chip.

### Co-culture of complex human and mouse microbiota on-chip

To determine whether the mouse Colon Chip can model symbiosis between microbiome bacteria and the intestinal epithelium, we seeded chips with healthy human microbiome (Hmb) or mouse microbiome (Mmb) that were previously shown to differ in their ability to promote intestinal barrier function both in gnotobiotic mice where these microbiota are maintained and passaged (2), and when cultured in human Colon Chips (14). When we compared the diversity between seeding stocks and the microbiome cultured on-chip for 40 hours, we found that on chip both microbiota exhibited similar alpha diversity (a metric of within sample taxonomic unit diversity) (**Supplementary Fig. S7**). However, the *Enterococcus* genus dominated the Hmb chips within 16 hours of colonization whereas Mmb chips contained multiple genera with variation in abundances between chips (**Fig. 3A**), which is likely due to the particulate nature of the dense microbiome sample when making dilutions and carrying out chip seeding. Compared to sterile chips without bacteria that were lined with nucleated epithelial cells covering from 97-100% of the surface area of the porous membrane, the presence of Hmb or Mmb induced variable levels of epithelial damage, with lesion areas covering from 9 to 94% of the membrane surface (**Fig. 3B**). Intestinal epithelial cell secretion of CXCL1 upon microbiome colonization also varied between chips, but CXCL1 secretion was not significantly different after 16 hours of colonization (**Fig. 3C**). These results indicate that the mouse Colon Chips can model a symbiotic relationship between healthy microbes and the colon epithelium; however, seeding a complex population of microbes, as is present within the Hmb and Mmb samples, results in variation in both microbiome composition and epithelial damage between chips.

### Colon Chips with microbiota can be used to select strains that protect against pathogen

We next asked whether the Colon Chips colonized with different microbiota can be used to identify particular bacterial strains that modulate host response to pathogens. Because the mouse microbiome protects against *S. typhimurium* infection in mice (2), we leveraged the variations in microbiome composition we observed between chips colonized with Hmb and Mmb to identify microbes that protect against *S. typhimurium* infection. When Colon Chips were seeded with Hmb or Mmb and infected with *S. typhimurium* 16 hours later, Hmb colonization inhibited *S. typhimurium-*induced CXCL1 release **(Fig. 3D)**. 16S sequencing revealed that *S. typhimurium* colonized both Hmb and Mmb chips (**Fig. 3E**). However, while *S. typhimurium* outgrew all of the other microbiome bacteria in Mmb chips, this pathogen did not overwhelm chips colonized by Hmb microbiome (**Fig. 3E**), which is dominated by *Enterococci* (**Fig. 3A,E**).

To assess whether a particular species of *Enterococcus* was responsible for protection against *S. typhimurium* overgrowth, we isolated *Enterococci* from our Hmb stock and identified a single *Enterococcus* species, *E. faecium* (**Supplementary Fig. S8**), that exhibited this protective phenotype. When we colonized mouse Colon Chips with either *E. faecium* alone, *S. typhimurium* alone, or initially seeded with *E. faecium* and then infected with *S. typhimurium* 16 hours later, both bacterial species grew in the chips, but when combined, neither species grew to the same level as monocolonized chips (**Fig. 4A**). The presence of *E. faecium* alone did not induce CXCL1 release and, in fact, it prevented CXCL1 release induced by *S. typhimurium* (**Fig. 4B**). When analyzed after 40 hours of colonization with *E. faecium*, there was a mild, but not statistically significant, increase in lesion formation (0-37% vs. 0-2% of epithelial surface area) when compared to sterile chips (**Fig. 4C,D, E**). In contrast, *S. typhimurium* infection caused detachment of large regions of the epithelium and induced a significant increase in epithelial lesion formation (14-96% lesion area) by 24 hours of infection (**Fig. 4C,D,E**). In chips colonized with *E. faecium*, the epithelium also displayed brighter tight junctions staining than *S. typhimurium* infected chips **(Fig. 4F)** suggesting that *E. faecium* colonization protects from significant tight junction damage. However tight junction outlines in sterile chips were significantly brighter than *E. faecium* colonized chips (**Fig. 4G**) suggesting that although *E. faecium* does not induce significant epithelial lesions, it does impact ZO-1 intensity. These findings are consistent with past studies that showed *E. faecium* promotes tolerance to *S. typhimurium* infection in mice (5,6), but that *E. faecium* overgrowth can also have harmful effects in hospitalized patients(22,23). Our results further suggest that *E. faecium* may prevent *S. typhimurium* overgrowth as well. These findings are consistent with past studies that showed *E. faecium* promotes tolerance to *S. typhimurium* infection in mice (5,6), and our results further suggest that *E. faecium* may prevent *S. typhimurium* overgrowth as well. These results show that mouse Colon Chips can be used to identify bacteria strains within complex microbiome that protect intestinal epithelium from pathogen overgrowth and that mouse colon chips are able to model the nuances of commensal pathobionts that have both protective and harmful effects.

## DISCUSSION

Our results demonstrate that mouse Intestine Chips can be created with cells derived from organoids isolated from duodenum, jejunum, ileum, or colons from wild type mice or transgenic Kaede mice that express GFP in all of their cells (15). We also showed that both complex species-specific microbiota or individual types of bacteria can be cultured in the Colon Chips and that bacterial-specific effects on epithelial cell adhesion, tight junctions, barrier function, mucus production, and cytokine release can be quantified directly. When challenged with pathogenic *S. typhimurium*, the mouse Colon Chips responded differently depending on whether the lumen was also colonized with normal human or mouse complex microbiome. In the course of these studies analyzing individual differences in the microbiome composition of chips, we identified *E. faecium* isolated from human microbiome as a bacterium that promotes host tolerance to infection as measured by prevention of epithelial cell detachment; however, the presence of this bacterium also induced small regions of epithelial detachment highlighting the potential of a single type commensal bacterium to produce both beneficial and injurious properties. Mouse Intestine Chips may therefore be useful for future mechanistic studies designed to pinpoint interactions between specific microbes and the intestinal epithelium.

The mouse Intestine Chips we described provide multiple novel experimental approaches that could be useful for future studies on host-microbiome interactions. The generation of Intestine Chips using organoids from Kaede mice opens up the possibility top track cell movement and divisions, in addition to enabling Intestine Chips to be created from mice with a variety of genetic backgrounds. This ability to generate Organ Chips with different genetic backgrounds can be especially helpful when comparing sterile chips to chips colonized with microbiome, as re-deriving germ free mice from different genetic backgrounds is difficult and expensive. We also showed that fluorescent bacteria can be imaged in live and fixed chips enabling future visualization and mechanistic studies focused on interactions between bacteria and host epithelium as well as among different bacteria cell populations. In addition, we were able to use the Colon Chips to recapitulate mouse intestinal injury and cytokine responses to infection with pathogenic *S. typhimurium* bacteria in vitro, and to reconstitute host-microbiome symbiosis by populating Colon Chips with complex living mouse and human microbiome. Finally, we showed that differences in the composition of complex microbiome cultured on-chip can be leveraged to identify specific bacterial strains, such as *E. faecium*, which modulate host tolerance to infection by *S. tymphimurium*. Taken together, these findings suggest that the mouse Colon Chip provides a modular system in which specific bacteria and epithelial cells from different regions of the intestine can be co-cultured to interrogate a wide variety of host-microbe interactions.

In our studies focused on identification of bacteria that promote tolerance to infection, mouse Colon Chips were infected with the intestinal pathogen, *S. typhimurium*, as a positive control. In mice, *S. typhimurium* infection causes bloody diarrhea, weight loss, inflammation in the small intestine, cecum, and colon, extrusion of epithelial cells, sepsis, and death(24–26). *S. typhimurium* grew well in the Colon Chips and induced similar epithelial damage (e.g., tight junction disruption, cell detachment) as well as increases in cytokines that recruit immune cells such as CXCL1, CXCL2, CCL20, and a non-coding RNA, AW112010, all of which closely mimic those previously observed in infected mice in vivo (19–21). These results indicate that the mouse Colon Chip can be used to investigate epithelium-pathogen interactions and that *S*.*typhimurium* can be used as a positive control for epithelial damage in these Organ Chip models.

To determine whether the mouse Colon Chip can model a symbiotic relationship between bacteria and the epithelium, we colonized mouse chips with healthy human microbiome (Hmb) or healthy mouse microbiome (Mmb) isolated from Hmb or Mmb mice(2). While we have previously shown that Hmb promotes barrier function in human chips(14), we found here that the mouse Colon Chips exhibited more variable responses to colonization with complex microbiome. Given the variation in the microbiome composition we observed between chips, these differences might account for the different phenotypic responses observed in this study. Future studies exploring different microbiome compositions, both in terms of bacterial strains or population diversity at the genus level, could be help to identify bacteria that promote epithelial integrity, which could have many applications in clinical settings and microbiome-based therapies. Interestingly, even though Hmb and Mmb stock solutions contained similar levels of alpha diversity and the Hmb stock was the same as we previously reported (14), *Enterococcus* came to dominate the Hmb mouse gut chips whereas Mmb chips contained a more diverse set of species. Taken together, these results indicate that mouse Intestine Chips could be used in the future to investigate colonization dynamics between certain microbes or microbial consortia and host species.

We decided to use the variation between chips to our advantage to investigate how the microbiome affects the host response *S. typhimurium* infection. Regardless of the starting microbiome composition, *S. typhimurium* outcompeted the species in Mmb chips. *S. typhimurium* also grew in Hmb chips, but *Enterococcus* prevented *S. typhimurium* from dominating the microbiome. These results appear to conflict with a previous finding that Mmb protects from *S. typhimurium* better than Hmb (2). However, that study used Swiss Webster (SW) mice while our Hmb, and Mmb samples were isolated from stool obtained from C57/BL6 mice because this is the strain we isolated our organoids from as well. Furthermore, SW Hmb mice had a dampened immune response compared to Mmb mice suggesting that the Hmb caused more severe *S. typhimurium* infection via its effect on the immune system, as opposed to Hmb bacteria affecting *S. typhimurium* pathogenicity without involving immune cells. In support of our findings, several studies have shown that *E. faecium* promotes tolerance to *S. typhimurium* infection in mice and *C. elegans* by an immune independent mechanism(5,6).

Because *Enterococcus* species such as *E. faecium* and *E. faecalis* can easily become resistant to vancomycin and cause sepsis in hospitalized patients, referred to as Vancomycin Resistant Enterococcus (VRE) disease, we sought to use the Colon Chips to: 1) determine if our *E. faecium* alone causes damage and 2) determine if our *E. faecium* can protect the epithelium from *S. typhimurium*-induced damage. Interestingly, we found that *E. faecium* protects from significant *S. typhimurium*-induced epithelial detachment, tight junction disruption, and CXCL1 induction, supporting findings in mice that *E. faecium* promotes a tolerance response to *S. typhimurium(5,6)* . However, colonization with *E. faecium* alone decreased tight junction intensity and appeared to produce some epithelial damage when analyzed across multiple chips, with lesions being observed in 0-37% of the epithelial surface in chips cultured with *E. faecium* alone vs. 0-2% in sterile chips and 14 to 96% in chips infected with *S. typhimurium*. Based on this observation, we suggest that *E. faecium* should not be used as a probiotic to prevent bacterial infections, and instead, understanding the bacterial molecules from *E. faecium* that protect against disease (5,6) and the mechanisms of action, might lead to more useful therapies for patients.

## Author Contributions

FSG designed and performed the experiments, analyzed data, and prepared the manuscript. DMC and MW analyzed 16S data. MP and FNG performed the experiments. AD generated *S. typhimurium mCherry*. MJC and MS discussed results. DLK designed experiments and discussed results. DEI designed experiments, discussed results, and prepared the manuscript with FSG.

## Acknowledgments

We would like to thank T. Ferrante with his assistance with imaging and L. Jin for her artwork.

