## Supplementary Figures and Methods for "Harnessing Colon Chip technology to identify commensal bacteria that promote host tolerance to infection"

#### *Supplementary Material*

#### SUPPLEMENTARY FIGURE LEGENDS

**Supplementary Figure S1. Cells isolated from organoids derived from different regions of small intestine and colon form confluent monolayers in microfluidic Organ Chips.** Organoids derived from duodenum, jejunum, ileum, and colon of C57/Bl6 mice were generated as previously described(16,17) and seeded onto microfluidic Organ Chips. Organoids from all parts of the intestine produced confluent monolayers within one week of seeding. Brightfield images of (**A**) duodenum (**B**) jejunum (**C**) ileum and (**D**) colon chips 13 days after seeding.

**Supplementary Figure S2. Colonic epithelial cells form a polarized monolayer in the mouse Colon Chip.** Movie shows Z- stack images taken every 2.25 microns of sterile chip from base of membrane towards top of apical channel. Nuclei (DAPI, blue) appear at the base followed by tight junctions (ZO-1, green) and mucus (MUC2, magenta) towards the top.

**Supplementary Figure S3. Muc2, ChrA, and Lgr5 gene expression in sterile Colon Chips relative to expression levels in colon organoids.** Sterile colon chips and organoids were maintained for 8 days before RNA extraction. *Muc2*, *ChrA*, and *Lgr5* RNA was detected in both chips and organoids, but *Muc2* and *ChrA* were expressed 98-fold and 498-fold higher, respectively in chips and *Lgr5* was expressed 10-fold higher in organoids.

**Supplementary Figure S4. *S. typhimurium* visualization on-chip.** Chips were infected with *S.typhimurium*-mCherry for 24 hours. *S. typhimurium* could be detected by **(A)** live microscopic imaging of *S. typhimurium* (magenta) and epithelium (green, Cell Tracker) as well as **(B)** after 4% PFA fixation (*S. typhimurium,* yellow; DAPI-stained nuclei (blue).

**Supplementary Figure S5. CXCL1 and CXCL2 are released 24 hours after S. typhimurium infection.** Chips were infected with *S. typhimurium* and basal outflow was collected 1.5, 3, 6, 24 hours after infection. (**A**) CXCL1 and (**B**) CXCL2 protein levels were increased 24 hours after infection whereas expression levels of other cytokines **(C)** were very low.

**Supplementary Figure S6. Gene expression 6 hours after S. typhimurium infection.** Mouse intestine chips were infected with *S. typhimurium* for 6 hours or maintained sterile and RNAs were analyzed by microarray sequencing; differentially expressed genes with a false discovery rate of $q<0.05$ are plotted.

**Supplementary Figure S7. Alpha diversity of Hmb and Mmb seeding stocks and chips.**

Observed alpha diversity (richness) and Shannon diversity of Hmb and Mmb stocks diluted 1:10, 1:1000, and after 40 hours on chip when seeded with 1:1000 dilution. There was a high degree of diversity within samples for microbiomes grown in the Colon Chips. When the chips were colonized with Hmb or Mmb for 16 hours, and then spiked with a human *E. coli* isolate for 24 hours, there was no significant impact on diversity.

**Supplementary Figure S8. Isolation and identification of *E. faecium* from the Hmb stock.** Hmb stock was plated on bile esculin agar (BEA) plates in which only *Enterococcus* species can hydrolyze bile esculin to produce black insoluble salts. 16S sequencing was performed on individual colonies picked from BEA plates. All sequences were identical and matched to *E. faecalis*, *E. faecium*, and *E. durans*. Growth in media containing L-Arabinose, Sorbitol, or Melibiose revealed acid production from L-Arabinose and Melibiose indicating *E. faecium*. Growth Vancomycin Resistant Enterococcus plates revealed purple colonies confirming Hmb *Enterococcus* is *E. faecium*.

#### SUPPLEMENTAL METHODS

##### RNA isolation, quantitative reverse transcription polymerase chain reaction, and microarray analysis

For RNA isolation, each channel of chips were washed with 100 ul of PBS. Cells were harvested using RLT buffer + BME and RNA was extracted using the Qiagen RNAeasy mini kit (74106, Qiagen). cDNA was synthesized using 0.5 ug RNA, 500 ng of random primers (Invitrogen 48190-011), 0.5 mM dNTPs, 1X First-Strand Buffer, 5mM DTT, and 100 U of Superscript III (Invitrogen 18080-044) according to the manufacturer's instructions. cDNA was diluted 1:4 with water and amplified using LightCycler 480 DNA SYBR Green I Master (Roche Applied Science 04887352001) and run on the LightCycler 96 (Roche 05815916001) using the preset Sybr Amp Melt Curve. The second derivative max was used to identify transcript copy number and normalized to the housekeeping gene, β-actin. The following primers were used: Muc2 F ctgaccaagagcgaacac, Muc2 R catgactggaagcaactgga TGCTGGGGTTTTTGTGAATCTC, ChrA F ATCCTCTCTATCCTGCGACAC, ChrA R GGGCTCTGGTTCTCAAACACT, Lgr5 F CCTACTCGAAGACTTACCCAGT, Lgr5 R GCATTGGGGTGAATGATAGCA, β-actin F GGCTGTATTCCCCTCCATCG, β-actin R CCAGTTGGTAACAATGCCATGT. For microarray RNA was extracted as described above and sent to Advanced Biomedical Laboratories for Microarray sequencing using the mouse Clariom D array. All analyses were run in R with custom scripts. Differentially expressed genes with a false discovery rate of $q<0.05$ were plotted.

##### Live Imaging of mouse colon chips infected with *S. typhimurium*

For live imaging of chips, Cell Tracker green was added to chips according to manufacturer's instructions (C2925 Thermo Fisher Scientific). Outlets and inlets were plugged with pipette tips that were cut to ¼ inch height and blocked with glue chips before imaging with a laser scanning confocal microscope (Leica SP5 X MP DMI-6000).

**Isolation of Enterococcus faecium from Hmb stock**

Hmb stock was plated on bile esculin agar (BEA) plates in which only Enterococcus species can hydrolize bile esculin to produce black insoluble salts. Colony PCR was performed on individual colonies. Individual colonies were inoculated into 20 ul of water and boiled at 95C for 10 minutes. 16S gene was amplified using RANGER mix (Bioline Bio-25052) according to the manufactur's instruction and the following 16S primers 27F: AGA GTT TGA TCM TGG CTC AG
1492R: CGG TTA CCT TGT TAC GAC TT and PCR cycle 94C 3 minutes followed by 35 cycles of:94C 45 second, 50C 60 seconds, 72C 90 seconds, ending with 72C for 10 minutes. PCR product was purified using Qiaquick PCR purification kit (Qiagen 28104) and sent to Dana Farber Sequencing Core for Sanger sequencing. All sequences were identical and matched to *E. faecalis*, *E. faecium*, and *E. durans*. Growth in media containing L-Arabinose (Anaerobe Systems AS 824), Sorbitol (AS 855), or Melibiose (AS 851) revealed acid production from L-Arabinose and Melibiose indicating *E. faecium*. Growth Vancomycin Resistant Enterococcus plates (R01830 Thermo Fisher Scientific) revealed purple colonies confirming Hmb *Enterococcus* is *E. faecium*.

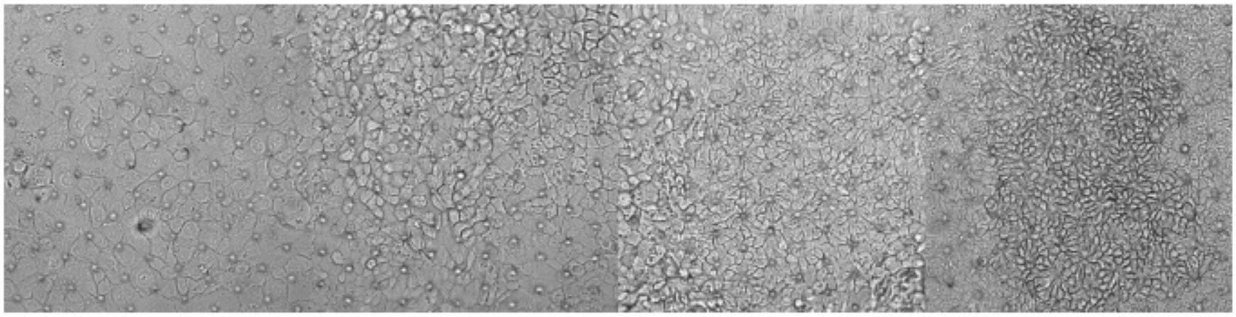

### Supplementary Figure S1

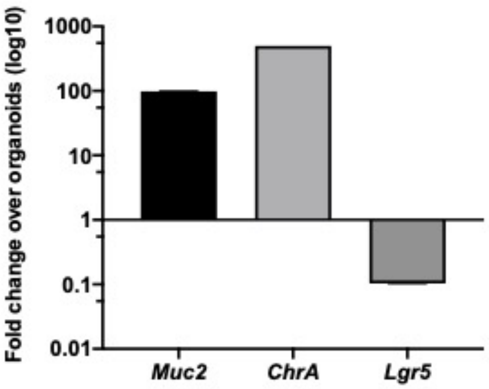

### Supplementary Figure S3

A

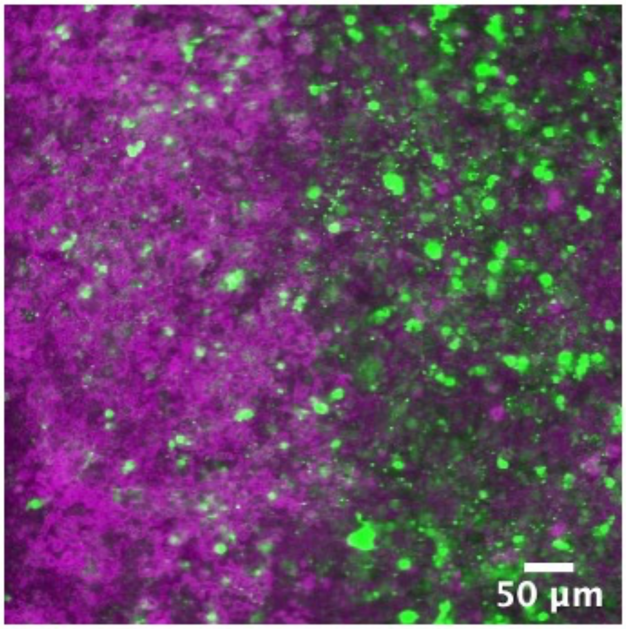

B

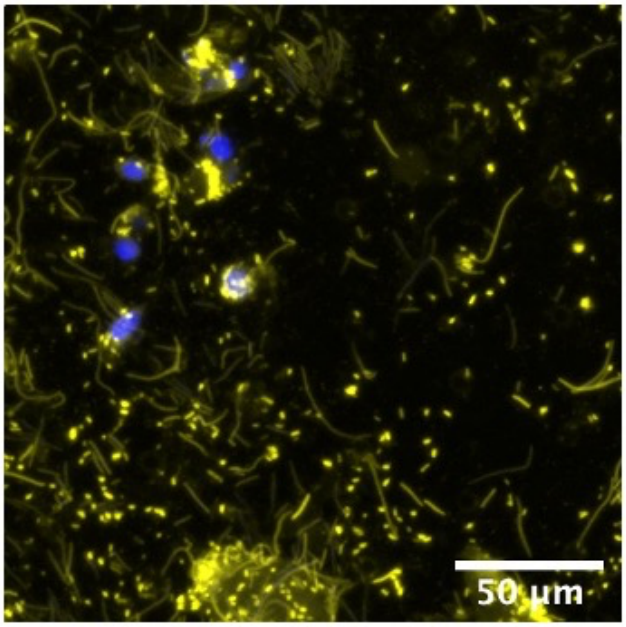

**Supplementary Figure S4**

A

B

C

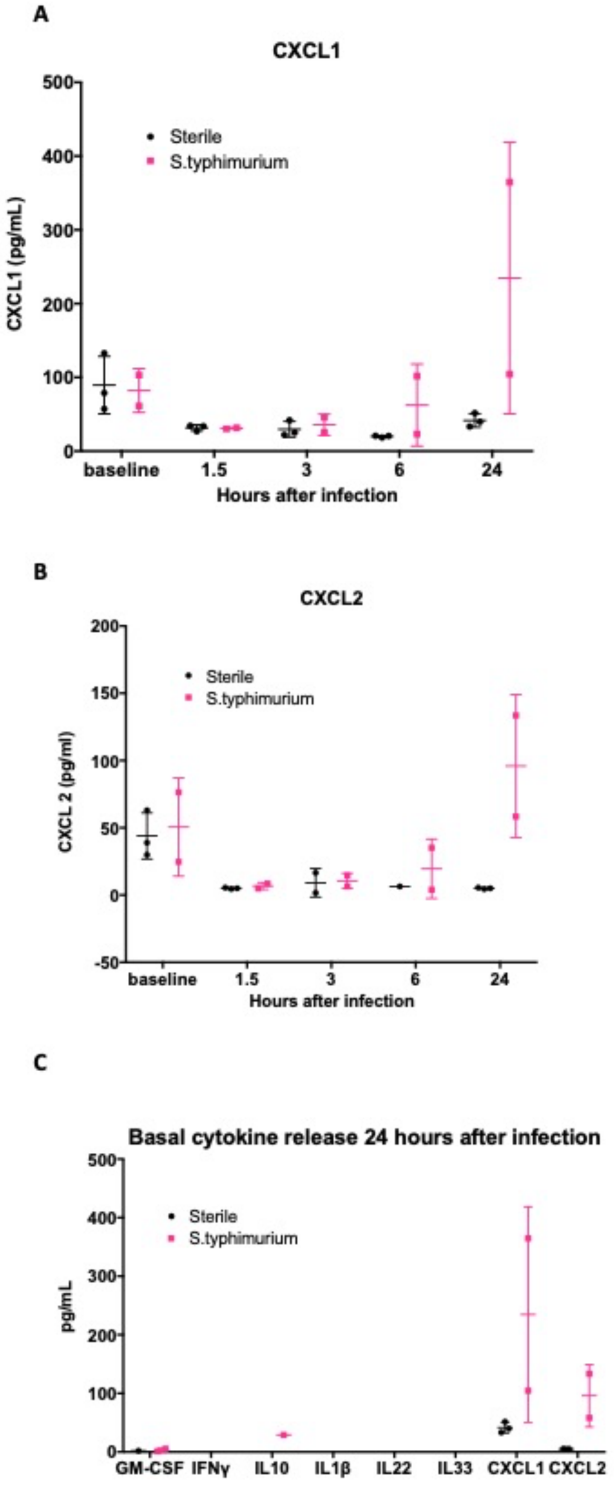

### Supplementary Figure S5

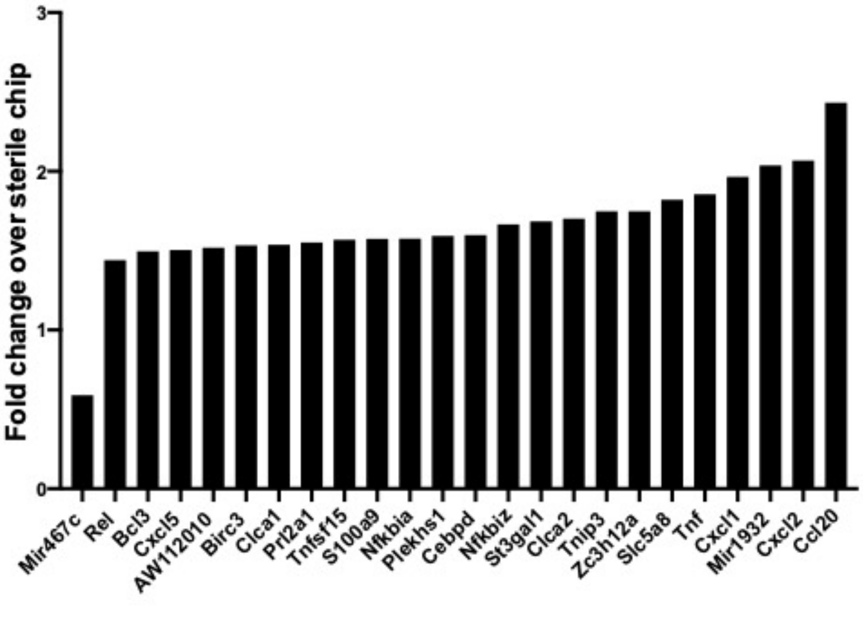

### Supplementary Figure S6

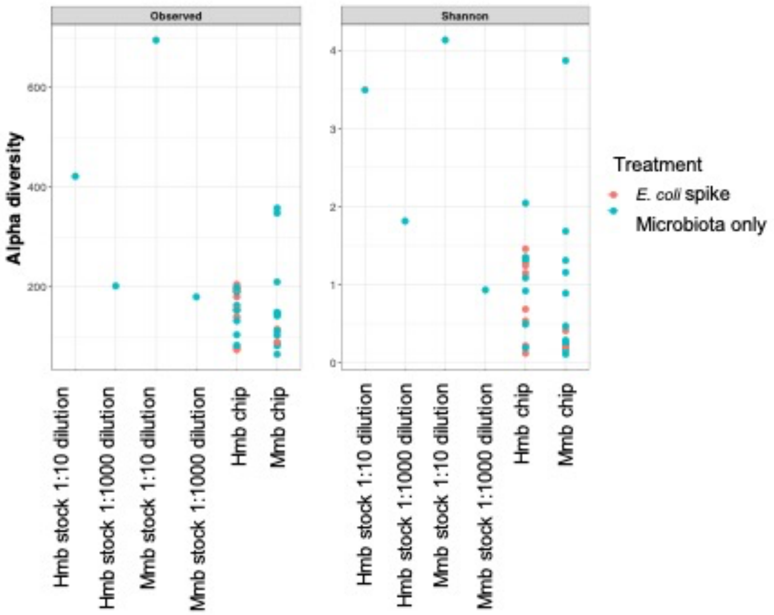

**Supplementary Fig. S7**

**1. Enterococcus selection**

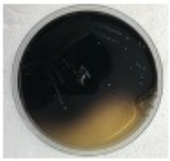

Plate Hmb stock on Bile esculin agar (BEA) plates. *Enterococcus* species hydrolize esculin to produce black insoluble iron salts

**2. Colony PCR**

Select colonies for 16s sequencing:
All isolates same16s sequence matches to:
*E. faecalis*
*E. faecium*
*E. durans*

**3. Biochemical tests**

| Acid from: | *E. faecalis* | *E. faecium* | *E. durans* | *Enterococcus* Hmb |
|---|---|---|---|---|
| L-Arabinose | | X | | X |
| Sorbitol | X | | | |
| Melibiose | | X | | X |

| VRE plate | *E. faecalis* | *E. faceium* | *Enterococcus* Hmb |
|---|---|---|---|
| Blue colonies | X | | |
| Purple colonies | | X | X |

### Supplementary Figure S8
